# Multi-Stain Fusion of Histopathology Images Using Deep Learning for Pediatric Brain Tumor Classification

**DOI:** 10.64898/2026.04.10.717785

**Authors:** Christoforos Spyretos, Iulian Emil Tampu, Per Nyman, Alia Shamikh, Joakim Lindblad, Neda Haj-Hosseini

## Abstract

The classification of pediatric brain tumors is investigated using deep learning on hematoxylin and eosin (H&E) and antigen Ki-67 (Ki-67) whole slide images (WSIs) from the Children’s Brain Tumor Network (CBTN) dataset. A total of 1,662 unregistered WSIs (1,047 H&E and 615 Ki-67 images) were analyzed, including low-grade glioma/astrocytoma (grades 1, 2) (LGG), highgrade glioma/astrocytoma (grades 3, 4) (HGG), medulloblastoma (MB), ependymoma (EP) and ganglioglioma. The The aim of this study was to effectively classify pediatric brain tumors using H&E and Ki-67 WSIs individually, and to investigate whether early, intermediate, and late fusion could improve the predictive performance. From each WSI, 224×224 pixel patches were extracted, and the instance (patch)-level features were obtained using the histology foundation model CONCHv1_5. The instances were aggregated using clustering-constrained attention multiple instance learning (CLAM) for patient-level classification. Model interpretability and explainability was assessed through attention heatmaps, cell density and Ki-67 labelling index (LI) maps. In the binary grade classification between LGG and HGG, the intermediate concatenation fusion achieved the best performance with a balanced accuracy of 0.88 ± 0.05, (*p* < 0.005) compared to the single-stain models (H&E: 0.84 ± 0.05, Ki-67: 0.86 ± 0.05). For the 5-class tumor type classification, the one-hidden layer late fusion learning model achieved the highest balanced accuracy of 0.83 ± 0.04 (*p* < 0.005), outperforming the single-stain models (H&E: 0.77 ± 0.05, Ki-67: 0.74 ± 0.05). Overall, most of the fusion approaches outperformed the single-stain models in both classification tasks (*p* < 0.005). The Ki-67 attention maps demonstrated moderate to strong Spearman correlation (*ρ* = 0.576 − 0.823) with the cell density and Ki-67 LI maps, suggesting that these features are associated with the model’s predictions, although additional features may contribute. The results show that H&E and Ki-67 images provide complementary information, and most of the multi-stain fusion approaches using deep learning improve pediatric brain tumor diagnosis.

## 1 Introduction

Central nervous system (CNS) tumors represented the second leading cancer among children and adolescents aged 0-19 years worldwide in 2022, with incidence and mortality rates of 1.2 and 0.6 per 100,000 individuals, respectively [1].CNS tumors include brain and spinal cord tumors, with this study focusing specifically on brain tumors. The diagnostic and prognostic assessment of brain tumors depends on the combination of clinical records, evaluation of non-invasive imaging modalities such as magnetic resonance imaging (MRI), and histopathological analysis of biopsy specimens. Additionally, molecular information is increasingly integrated into clinical practice, as reflected in the fifth edition of the World Health Organization (WHO) classification guidelines for CNS tumors, published in 2025 [2].Despite substantial advances in molecular diagnostics, molecular profiling is costly, time-consuming and not widely available, with histological assessment continuing to be essential for evaluating brain tumors and guiding the selection of appropriate ancillary molecular tests. Histological examination is labor-intensive, and the limited availability of expert neuropathologists in pediatric brain tumors emphasizes the importance of decision-support tools to aid diagnosis [3].The digitization of histology glass slides into whole slide images (WSIs) and the growing availability of large WSI datasets has enabled the development of computational pathology (CPath). Within CPath, the implementation of deep learning has been facilitated, with the goal of optimizing and assisting healthcare professionals’ workflows, leading to better and more effective patient care. Deep learning models applied to WSIs have been used for a wide range of tasks, including tissue classification (e.g., distinguishing benign from malignant tissue, subtyping, and grading), detection (e.g., lymphocyte or mitosis counting), segmentation (e.g., nuclei and gland delineation), prediction of molecular markers, and patient survival and prognostic modeling [4].

Early success in deep learning for brain tumor WSI classification largely relied on supervised learning frameworks, in which models were trained on fully labeled datasets, typically using patches extracted from WSIs along with their corresponding patch-level annotations. However, obtaining a large number of images with finegrained annotations at a patch level is labor intensive, costly, and time consuming for pathologists to annotate them. In addition, annotation labels could often subjective and of limited accuracy. To address this, weakly and self-supervised learning methods have been developed to perform slide-level classification. Under the weakly-supervised learning paradigm, models are trained on partially or sparsely labeled data, such as one label for an entire WSI or per patient. With self-supervised learning, the algorithms learn feature representations through unlabeled data. Multiple instance learning (MIL) is a weakly supervised learning method, which has been widely used in WSI classification tasks with recognized success [5, 6].In MIL each WSI is considered as a bag containing multiple patches, also called instances. If a WSI is labeled class-positive, then at least one patch in that WSI is class-positive. Otherwise, if a WSI is class-negative, all patches in that WSI are negative [7].Several reviews provide thorough comparisons of MIL approaches in CPath, highlighting their limitations, challenges, and future potential [8, 9, 10, 11].The recent emergence of histology foundation models has transformed the field of CPath. Histology foundation models are trained on immense histology datasets, capturing a broad spectrum of patterns such as different fixations, staining characteristics, scanning protocols and tissue architecture across different centers, compared to models trained on out-of-domain datasets, such as ImageNet. Histology foundation models could be effectively utilized as feature extractors for various slide and patient-level classification tasks, mitigating generalization and domain shift challenges, and are label efficient for zero-shot and few-shot slide classification for rare and underrepresented tumors [12].

The increasing availability of medical data in CPath, such as medical images, electronic health records and genome sequences, has led to the development of multimodal deep learning approaches [13, 14].These approaches aim to improve predictive performance by integrating data across diverse modalities, mimicking the multimodal nature of clinical practice. In the context of histology and pediatric brain tumors, pathologists often use various immunohistochemical (IHC) stains in addition to hematoxylin and eosin (H&E) to identify specific molecular alterations, detect mutant proteins, determine molecular subgroups and malignancy [15, 16].One such IHC marker is the Antigen Kiel 67 protein (Ki-67), which is known for staining proliferating (positive) and non-proliferating (negative) cells. In this regard, the ratio of the number of the proliferating cells to the total number of cells, known as the Ki-67 labeling index (LI), is a key component in assessing tumors [17, 18].An increased Ki-67 LI is associated with higher malignancy in gliomas and is used to differentiate between low-grade and high-grade gliomas [19, 20].In adult medulloblastoma, Ki-67 labeling index (LI) is correlated with molecular subgroups and poorer overall survival [21].In the pediatric population, the grades and prognosis of ependymoma are correlated with Ki-67 LI [22].Additionally, in pediatric medulloblastoma, the Ki-67 LI have a potential prognostic use [23], with high Ki-67 LI observed in anaplastic tumors, whereas non-anaplastic tumors exhibit lower values [24].The examination of Ki-67 WSIs could be specifically valuable in low-resource settings, in which the molecular examination is inaccessible. To the best of our knowledge, no study has investigated the fusion of registered or unregistered H&E and Ki-67 images using deep learning for pediatric or brain tumor diagnosis, and such approaches remain limited across other cancer types, leaving the potential of multi-stain fusion with deep learning in CPath unexplored.

In this study, it is investigated if pediatric brain tumors could be effectively classified using only H&E or Ki-67 slides. It is further examined whether the fusion of unregistered H&E and Ki-67 slides could improve the predictive performance of pediatric brain tumor families/types compared to only using a single stain modality. Moreover, the explainability and interpretability of the Ki-67 attention maps are quantitatively assessed using the Ki-67 LI maps, and the negative and positive cell density maps.

## 2 Background and Related Work

A limited number of studies have applied deep learning to pediatric brain tumors for classification or survival prediction [25, 26, 27, 28, 29], with research primarily focused on adult brain tumors. In [25], a deep learning method has been proposed for survival prediction in pediatric and adult brain tumors using H&E WSIs and genetic data, with slide-level representations obtained by averaging patch-level features extracted using an ImageNet-pretrained ResNet50 model. Three multimodal fusion strategies were evaluated, and the results indicated that models trained only on H&E WSIs achieved a mean composite score of 0.854 for pediatric brain tumor survival prediction, which improved by 0.065 when H&E features were early fused with RNA data. In medulloblastoma pediatric brain tumors, CNN-based approaches demonstrated high accuracy for classifying classic, desmoplastic, large cell, and nodular subtypes through late fusion of H&E features with textual information [27], while an ImageNet-pretrained Efficient-Net model achieved a patch-level F1-score of 0.801 for distinguishing classic and desmoplastic/nodular subtypes [28].In [29], a linear discriminant analysis (LDA) model was trained on features extracted from pathologist-selected regions of interest (ROIs) in WSIs to classify SHH and WNT-activated versus Group 3 and 4 medulloblastoma subtypes, providing a patient-level area under the receiver operating characteristic curve (AUROC) of 0.70, and subsequent survival prediction within each subtype, with the highest AUROC of 0.92 obtained for Group 3 medulloblastoma. In our previous work [26], a hierarchical classification of pediatric brain tumors was performed using the UNI histology foundation model to extract features from H&E WSIs, combined with the attention-based MIL (ABMIL) framework. The hierarchy progressed from broad tumor categories (seven classes) to families (10 classes) and types (nine classes). The model reached Matthew’s correlation coefficients (MCC) of 0.76 ± 0.04, 0.63 ± 0.04, and 0.60 ± 0.05 for tumor category, family, and type classification, respectively.

The fusion of multiple stains has been explored only to a limited extent in cancers of other organs [30, 31].In [30], a multimodal pretraining strategy aligning H&E and IHC (ER, PR, HER2, Ki67) WSIs demonstrated improved performance across WSI classification, molecular subtyping, and prognostic prediction in breast cancer and kidney transplants. A knowledge transferred from a foundation model pretrained on H&E WSIs to IHC WSIs was investigated in [31], providing better results in H&E patch-level colorectal cancer subtyping and WSI classification, and IHC marker expression prediction. These studies suggested that fusing stain information can improve performance, leaving the potential of fusing H&E with IHC and other stain WSIs in deep learning pediatric brain tumor analysis to be further investigated.

## 3 Dataset

In this study, the dataset was obtained from the Children’s Brain Tumor Network (CBTN), consisting of *∼* 2,000 subjects and *∼* 8,000 multiple stained slides collected from 32 institutions across the United States, Australia, Switzerland, and Italy [32, 33].The WSIs have a *×* 20 magnification with a pixel size ranging between 0.251 and 0.505 *µ*m. The slides are unregistered and many subjects contain multiple single-stained WSIs, with H&E WSIs being the most common and Ki-67 being one of the most representative IHC staining protocols. Thus, only subjects with slides containing the two stains were selected to conduct the experiments. The dataset is based on pre-2021 WHO classification of tumors of the CNS guidelines, in which some tumor entities are no longer used in the current clinical practice. Therefore, in accordance with clinicians, subjects with outdated classifications were excluded, and the classifications defined in the 2021 WHO guidelines were used to conduct the experiments [2].The quality of the WSIs were visually assessed by at least 1 nonpathologist, and selected images were revised by at least 1 pathologist. Based on the pathologists’ guidance, WSIs with light exposure, out-of-focus images, or originating from non-brain tissue (e.g., liver) were not included in the experiments. Additionally, WSIs with artifacts, such as pen marks and air bubbles, were included in the experiments, as these artifacts were rare and their effect on the model’s explainability has previously been shown to be negligible on the same dataset [34].

A threshold of at least 10 subjects per class was set, ensuring a sufficient representation of each brain tumor. Consequently, the final cohort consisted of 529 subjects (males/females: 290/239, mean age in years: 10.20 ± 5.93, age in years range: [0.15, 36.50]) and a total of 1,662 WSIs, in which 1,047 are H&E and 615 are Ki-67 WSIs. This study analyzed the tumor families of low-grade glioma/astrocytoma (grades 1, 2) (LGG) and high-grade glioma/astrocytoma (grades 3, 4) (HGG), and the tumor types of medulloblastoma (MB), ependymoma (EP) and ganglioglioma (GG). These entities follow the 2021 WHO hierarchical brain tumor classification, which progresses from broad tumor categories to families and types. In this study, all entities are referred to as tumor types for simplicity. Figure 1 illustrates representative H&E and Ki-67 WSIs with their close-up regions, and Table 1 summarizes the number of subjects and WSIs included in the study. The Ki-67 LI values for each tumor were calculated in [35].

**Table 1.** Tumor types, the number of subjects, and H&E and Ki-67 WSIs used in the analysis. Only subjects having WSIs from both the H&E and Ki-67 stain modalities were included in the experiments.

| Tumor type | Number of Subjects | Number of H&E WSIs | Number of Ki-67 WSIs |
| --- | --- | --- | --- |
| Low Grade Glioma (LGG) | 249 | 482 | 285 |
| High Grade Glioma (HGG) | 85 | 170 | 97 |
| Medulloblastoma (MB) | 76 | 127 | 82 |
| Ependymoma (EP) | 71 | 153 | 89 |
| Ganglioglioma (GG) | 48 | 115 | 62 |
| Total | 529 | 1047 | 615 |

**Figure 1.**
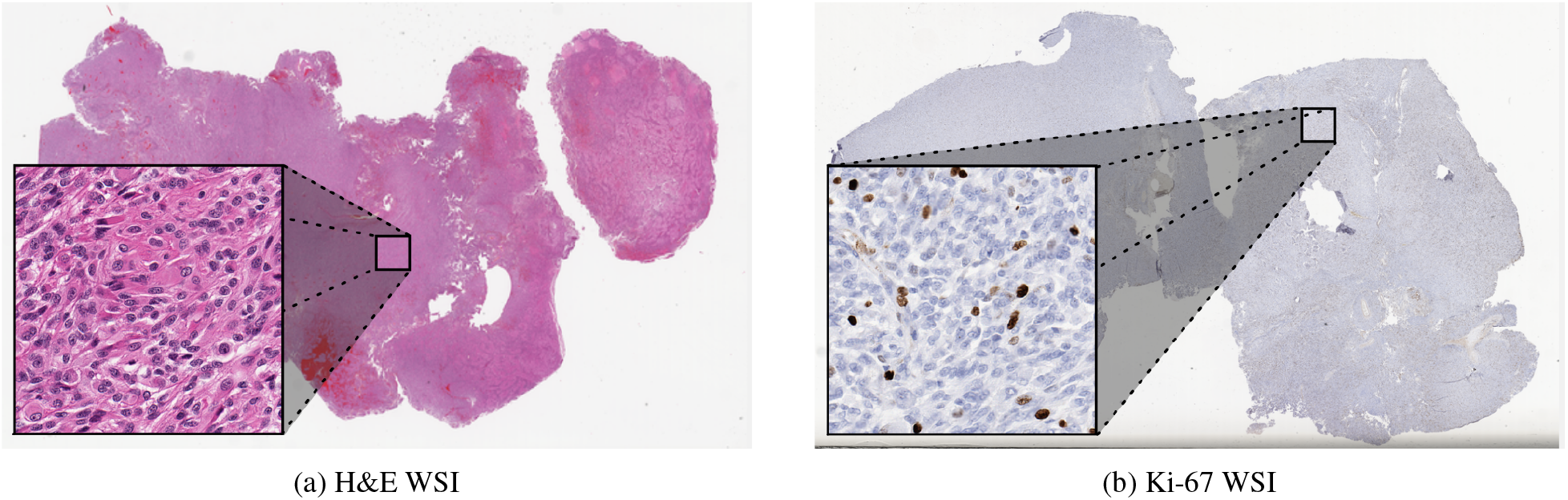
Representative (a) H&E and (b) Ki-67 HGG WSIs from the same subject. In the Ki-67 WSI the brown cells are proliferating (positive) cells and the blue cells are non-proliferating (negative) cell.

## 4 Methodology

The clustering-constrained attention multiple instance learning (CLAM), a MIL framework, was utilized to perform patient-level classification [36].First, patches and their corresponding features were extracted from each WSI. Subsequently, independent experiments were conducted for the H&E and Ki-67 WSIs, and fusion approaches between the stains were investigated with the aim to improve predictive performance. A schematic of the overall methodology is illustrated in Figure 2.

**Figure 2.**
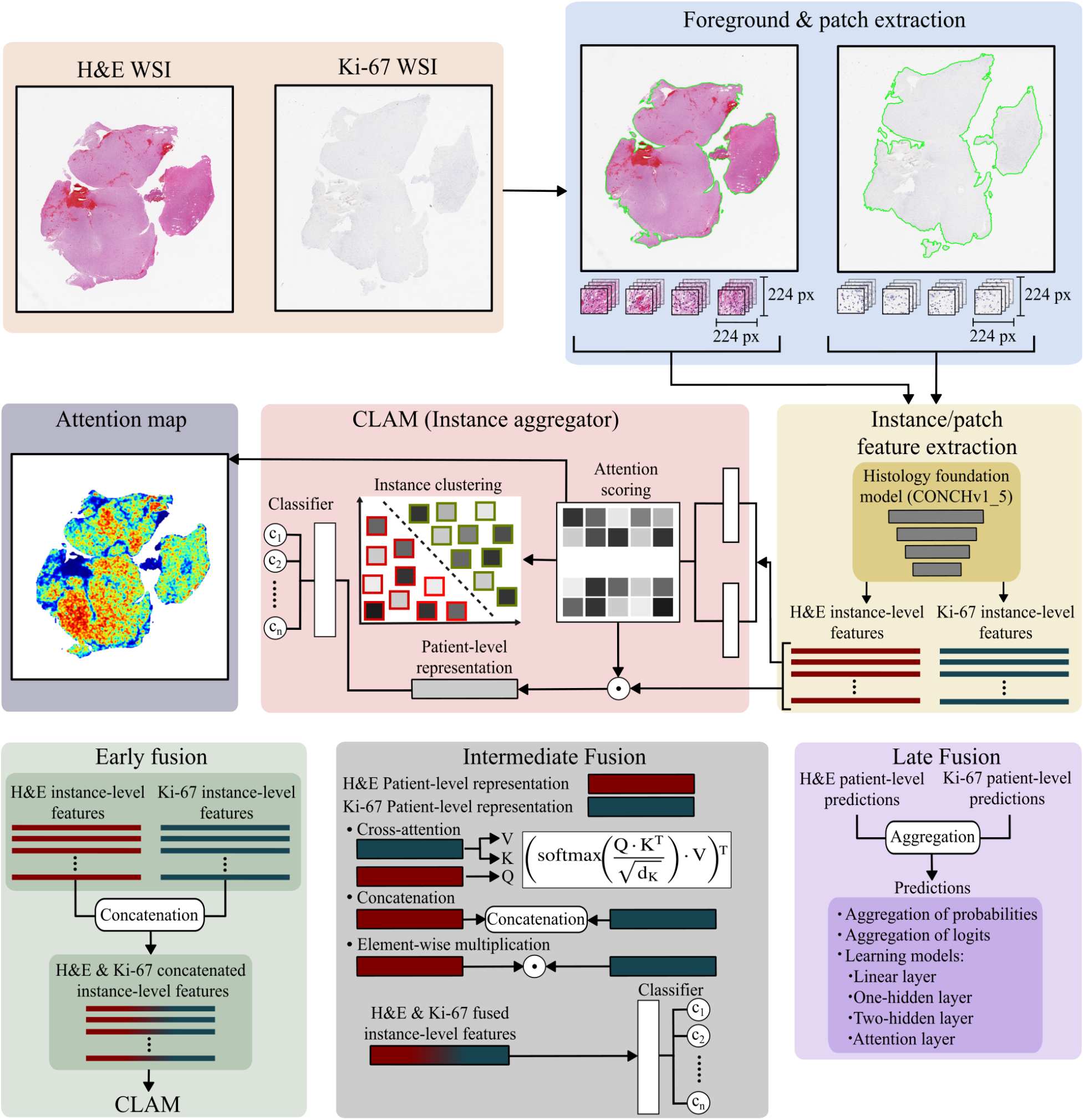
Overview of the methodology. Foreground and 224 *×* 224 patch extraction were performed on the WSIs, followed by instance/patch-level feature extraction using the CONCHv1_5 histology foundation model. Patient-level classification was conducted using CLAM separately on H&E and Ki-67 WSIs. Attention maps were generated to interpret the models’ predictions. Early, intermediate and late fusion approaches were explored to combine the information between the two stains.

### 4.1 Patch Extraction

The tissue regions were segmented and the background was excluded from the WSIs using the CLAM toolbox with default settings, followed by the extraction of nonoverlapping 224 *×* 224-pixel tissue patches (instances). The results were visually inspected, and settings were adjusted for slides in which tissue segmentation was not satisfactory.

### 4.2 Feature Extraction

The patch-level feature extraction was performed using the CLAM toolbox and the histology foundation model CONCHv1_5 [37].CONCHv1_5 is a large vision transformer model initialized from the UNI checkpoint [38] and fine-tuned on 1.17 image-caption pairs in histopathology following a procedure similar to that used for CONCH [39], with the dimensionality of the extracted features being 768. After extracting the patch-level features, those corresponding to the same stain modality for each subject were concatenated along the patch axis, representing patient-level features of shape number of patches *×* feature dimensionality per stain modality.

### 4.3 Single-Stain

CLAM uses a learnable pooling mechanism, which is based on the attention framework but is invariant to the number and order of instances within a bag. Two fully connected layers are used to assign a weight to each instance, and the bag-level representation is computed as the weighted average of the instances. A clustering layer is incorporated parallel to the attention mechanism and is trained to differentiate between instances that contribute positively or negatively to the correct patient label. The instance-level clustering loss is applied to direct the model’s attention toward instances that are informative of the true label, with a predefined subset of instances with the highest and lowest attention scores selected for clustering. Each patient representation is summarized as a single attention vector, computed as the product of the attention weight matrix and the patch-level feature embeddings, and then input to a linear classifier to provide the patient-level prediction. The CLAM model was trained and evaluated separately for the H&E and Ki-67 WSIs.

### 4.4 Multi-Stain Fusion

Varieties of early, intermediate and late fusion approaches were implemented to integrate complementary information across the stain modalities with the goal to improve the predictive performance [40].

#### 4.4.1 Early Fusion

In early fusion, information from all modalities are integrated at the input level before being fed into a single model. The modalities can be represented as raw data, hand crafted, or deep features, with the fused representation built through operations such as vector concatenation, elementwise summation and multiplication. With this approach, only a single model is trained, simplifying the design of the process. In this study, the patch-level features extracted from the H&E and Ki-67 WSIs of the same subject were concatenated along the patch axis to form patient-level features. Subsequently, the concatenated features were used to train and evaluate the CLAM model for patient-level classification.

#### 4.4.2 Intermediate Fusion

In intermediate fusion, modality-specific feature representations are iteratively refined through cross-modal information sharing during training. However, in this work, conventional intermediate fusion approaches could not be employed since features were extracted using the pretrained CONCHv1_5 histology foundation model, with feature extraction layers being frozen and not updateable during training. Therefore, the patient-level attention vectors were obtained during the training, validation, and test phases of the single-stain experiments. These vectors summarize the most informative features from all patches of the corresponding stain modality. Cross-attention, concatenation and element-wise multiplication were applied to fuse the H&E and Ki-67 patient-level attention vectors.

In the cross-attention, one stain modality was used as the query while the other modality was used as the keys and values [41, 42].This approach enables mutual information exchange between modalities, allowing each stain to inform the other. Let *M*_*i*_ *∈* R^1*×d*^ and *M*_*j*_ *∈* R^1*×d*^ denote the patient-level vectors for stain *i* and stain *j*, respectively, where *i, j ∈ {* H&E, Ki-67*}* and i ≠*j*. Since these are single vectors rather than sequences, the projected vectors were transposed from R^1*×d*^ to R^*d×*1^ so that the attention operates across the feature dimensions, resulting in a *d × d* attention matrix. Without the transposition, the attention matrix would be a scalar, and the softmax of a single value is equal to 1. The cross-attention was computed as follows:

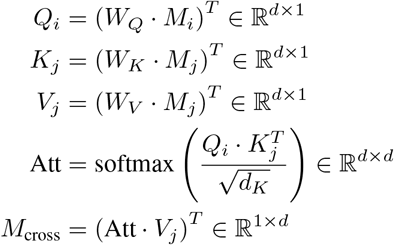

where *W*_*Q*_, *W*_*K*_, and *W*_*V*_ are learnable projection matrices for the query, key, and value, respectively, *d*_*K*_ is the dimension of the key vector, and *M*_cross_ is the cross-attended patient vector informed by the complementary stain modality. In the intermediate H&E-guided cross-attention fusion, Ki-67 is informed by H&E, whereas in the intermediate Ki-67-guided cross-attention fusion, H&E is informed by Ki-67.

In the concatenation approach, the H&E and Ki-67 patient-level vectors were concatenated along the feature dimension, resulting in fused vectors of size 1 *×* 2*d*. For the element-wise multiplication, the element-wise product was computed between the H&E and Ki-67 patient-level vectors. For each intermediate fusion approach, the resulting fused patient single vectors were used to train and evaluate a linear classifier, the same as in the CLAM framework.

#### 4.4.3 Late Fusion

In late fusion, a separate model is trained for each modality and the predictions from the individual models are aggregated to obtain a final prediction. With this approach, modality-specific architectures could be implemented. Late fusion is effective in the presence of missing, limited, or incomplete modalities, as each model is trained independently, and new modalities could be incorporated without retraining the entire model. Late fusion is often preferred due to its smaller number of parameters compared with early and intermediate fusion strategies. Moreover, since the errors of different unimodal models are often uncorrelated, late fusion approaches potentially results in reduced bias and variance in the aggregated predictions. Finally, the risk that modalities with higher information density disproportionately dominate the prediction is mitigated by allowing equal or modality-specific weighting during the aggregation step.

Three late fusion approaches were conducted, aggregation of softmax scores, aggregation of logits and using four learning models. The learning models included a linear model mapping directly to the output classes, a model with one-hidden layer, a model with two-hidden layers, and a model using an attention mechanism layer. The learning models were trained using the patient-level logits of the single-stain models as input features.

### 4.5 Experimental Setup

The classification tasks included binary grade classification between LGG versus HGG, and multi-class classification across the five tumors types. The dataset was class-stratified and patient-wise split into 70% for training, 10% for validation, and 20% for testing to prevent data leakage across the sets [43].The splitting procedure was repeated using non-parametric bootstrapping with 50 replicates, resulting in 50 independent train, validation and test splits. Non-parametric bootstrapping involves repeatedly sampling from the dataset with replacement to generate multiple training, validation, and test sets (replicates), allowing to estimate statistical measures without making data distribution assumptions.

### 4.6 Model Training Parameters

#### 4.6.1 Single-Stain

The classification tasks were performed using the small version of the single-branch CLAM model with the attention net gated mechanism. Models were trained using the AdamW optimizer, with weight decay of 0.00001, and a starting learning rate of 0.0001, which is decreased during training using a cosine annealing scheduler. The Adamw optimizer and the cosine annealing scheduler were additionally added to the CLAM original framework. For regularization, the dropout rate was set at 0.25 and applied after the initial fully-connected layer that processes input embeddings. Training was supervised by a patient-level cross-entropy loss (bag loss) and an instance-level smooth-SVM loss, with the parameter controlling the contribution of the two losses set to its default value of 0.7. To address class imbalance, the class-weighting adjustment was enabled for the patient-level cross-entropy loss. Furthermore, the default value of 8 was used for the number of high and low-attention instances used for the instance-level clustering. The number of epochs was set at 50, and early stopping was utilized on the cross-entropy validation loss with minimum number of epochs set at 10, patience at 5 epochs and stopping epoch at 20 epochs. All other settings were set at the default values suggested by [36].

#### 4.6.2 Early Fusion

The training of the early fusion models was performed using the same parameters as those employed for the single-stain models.

#### 4.6.3 Intermediate Fusion

The same training parameters as the single-stain models were used to train and validate the classifier, with the exception of the learning rate, which was set at 0.001.

#### 4.6.4 Late Fusion

Different training parameters for the learning models were defined based on the classification tasks and the model architecture. All models were trained using the Adam optimizer with a learning rate of 0.001 and a weight decay of 0.0001. Class weights were applied in the crossentropy loss to account for imbalanced data, and L1 regularization was used with a lambda value of 0.001.

In the LGG and HGG binary classification task, the single-layer and attention models were trained for 1500 epochs with early stopping patience set to 200 epochs. The one-hidden-layer and two-hidden-layer models were trained for 1200 epochs with early stopping patience of 100 epochs, with the one-hidden-layer model using a hidden dimension of 4 and the two-hidden-layer model employing hidden dimensions of 6 and 2 in the first and second layers, respectively.

In the 5-class classification task, the single-layer and attention models were trained for 1500 epochs, with early stopping patience set to 200 epochs for the single-layer model and 30 epochs for the attention model. The onehidden-layer and two-hidden-layer models were trained for 1200 epochs with early stopping patience of 150 epochs, with the one-hidden-layer model using a hidden dimension of 15 and the two-hidden-layer model employing hidden dimensions of 15 and 10 in the first and second layers, respectively.

### 4.7 Evaluation and Statistical Analysis

Since the class distribution of the dataset is imbalanced, the models’ performance was assessed using balanced accuracy, MCC, area under the receiver operating characteristic curve (AUC-ROC), weighted F1-score and mean confusion matrices. Balanced accuracy was chosen as the primary metric because it is suitable for both binary and multiclass imbalanced classification, whereas MCC is primarily used for binary classification. For the statistical comparison, two-sided permutation tests with 10,000 permutations were conducted on the test sets to assess differences in the performance of the models. Permuatation test is a non-parametric test and does not rely on distributional assumptions. Statistical comparisons were conducted between the single-stain and the fused stain modalities models, as well as comparisons between the fused models. Notably, only the best-performing late fusion and the best-performing intermediate fusion approach were included in the comparisons. The significance level was set to *α* = 0.05, and Bonferroni correction was applied to adjust the significance level when more than one comparison was performed.

### 4.8 Interpretability and Explainability

Utilizing the CLAM toolbox, attention maps were generated by the single-stain models to visualize and interpret the importance of different regions within the WSIs. The Ki-67 LI, negative and positive cell density maps were extracted using the method proposed in [35], with Ki-67 LI maps being computed as the ratio of tumor proliferating cells to the total cells. The attention heatmaps and LI, negative and positive cell density maps, which were normalized during generation, were resized to have the same spatial alignment and foreground (tissue) pixels. The Spearman’s rank correlation coefficient was computed to quantify the relationship between the pixel intensities of the Ki-67 attention heatmaps and the corresponding LI, negative and positive cell density maps. Additionally, the Ki-67 attention heatmaps of a few cases were visually compared with the corresponding Ki-67 LI, negative and positive cell density maps to assess whether regions assigned high attention by the model overlap with tumor areas exhibiting high and low proliferation.

### 4.9 Hardware and Software Environment

All the model trainings and evaluations were conducted on a computer equipped with an NVIDIA GeForce RTX 5090 GPU with 32 GB of memory, a 12-core CPU and a 128GB RAM. The codebase was implemented in Python (v3.12.0) using PyTorch (v2.9.1) with CUDA 13.0 support.

## Results

### 5.1 LGG vs HGG Classification Performance

The results of the binary grade classification are depicted in Table 2 and Figures 3 and 4. The Ki-67 single-stain model outperformed the H&E single-stain model across all evaluation metrics, and the statistical comparison at a significance level of *α* = 0.05 demonstrated significant differences in balanced accuracy, MCC, and weighted F1-score (*p <* 0.05), while no significant difference was observed for AUC-ROC (*p >* 0.05).

**Table 2.** Binary classification performance between LGG and HGG on the test sets. Metrics are reported as a mean ± standard deviation with 95% confidence intervals (CI) shown in brackets, computed across 50 replicates of nonparametric bootstrapping. The best performing approach and metrics are highlighted in bold.

| Model | Balanced Accuracy | MCC | AUC-ROC | Weighted F1-score |
| --- | --- | --- | --- | --- |
| Single-Stain H&E | 0.84 ± 0.05<br>[0.82, 0.85] | 0.70 ± 0.09<br>[0.67, 0.72] | 0.91 ± 0.04<br>[0.90, 0.92] | 0.88 ± 0.03<br>[0.87, 0.89] |
| Single-Stain Ki-67 | 0.86 ± 0.05<br>[0.84, 0.87] | 0.74 ± 0.09<br>[0.72, 0.77] | 0.92 ± 0.05<br>[0.91, 0.93] | 0.90 ± 0.03<br>[0.89, 0.91] |
| Early Fusion | 0.87 ± 0.05<br>[0.86, 0.88] | 0.76 ± 0.08<br>[0.73, 0.78] | **0.94 ± 0.04**<br>**[0.93, 0.95]** | **0.91 ± 0.03**<br>**[0.90, 0.91]** |
| Intermediate H&E-Guided Cross-Attention Fusion | 0.87 ± 0.05<br>[0.85, 0.88] | 0.73 ± 0.09<br>[0.71, 0.76] | 0.92 ± 0.04<br>[0.91, 0.93] | 0.90 ± 0.03<br>[0.89, 0.91] |
| Intermediate Ki-67-Guided Cross-Attention Fusion | 0.86 ± 0.06<br>[0.85, 0.88] | 0.73 ± 0.09<br>[0.70, 0.76] | 0.92 ± 0.04<br>[0.91, 0.93] | 0.90 ± 0.04<br>[0.89, 0.91] |
| **Intermediate Concatenation Fusion** | **0.88 ± 0.05**<br>**[0.86, 0.89]** | 0.76 ± 0.09<br>[0.74, 0.79] | 0.92 ± 0.04<br>[0.91, 0.94] | **0.91 ± 0.03**<br>**[0.90, 0.92]** |
| Intermediate Element-Wise Multiplication Fusion | 0.87 ± 0.05<br>[0.85, 0.88] | 0.75 ± 0.09<br>[0.73, 0.78] | 0.93 ± 0.04<br>[0.92, 0.94] | **0.91 ± 0.03**<br>**[0.90, 0.92]** |
| Aggregation of Softmax Scores Late Fusion | 0.87 ± 0.05<br>[0.86, 0.88] | **0.77 ± 0.08**<br>**[0.75, 0.79]** | 0.93 ± 0.04<br>[0.92, 0.94] | **0.91 ± 0.03**<br>**[0.90, 0.92]** |
| Aggregation of Logits Late Fusion | 0.87 ± 0.05<br>[0.86, 0.88] | **0.77 ± 0.08**<br>**[0.75, 0.79]** | 0.93 ± 0.04<br>[0.92, 0.94] | **0.91 ± 0.03**<br>**[0.90, 0.92]** |
| Linear Layer Late Fusion Learning Model | 0.83 ± 0.12<br>[0.80, 0.86] | 0.68 ± 0.24<br>[0.61, 0.74] | 0.91 ± 0.08<br>[0.89, 0.93] | 0.88 ± 0.09<br>[0.85, 0.90] |
| One-Hidden Layer Late Fusion Learning Model | 0.87 ± 0.07<br>[0.85, 0.89] | 0.75 ± 0.14<br>[0.71, 0.78] | 0.92 ± 0.04<br>[0.91, 0.93] | 0.90 ± 0.05<br>[0.89, 0.92] |
| Two-Hidden Layer Late Fusion Learning Model | 0.87 ± 0.07<br>[0.85, 0.89] | 0.74 ± 0.14<br>[0.70, 0.78] | 0.92 ± 0.04<br>[0.91, 0.93] | 0.90 ± 0.05<br>[0.88, 0.91] |
| Attention Layer Late Fusion Learning Model | 0.78 ± 0.18<br>[0.74, 0.83] | 0.56 ± 0.34<br>[0.46, 0.65] | 0.88 ± 0.15<br>[0.84, 0.92] | 0.82 ± 0.17<br>[0.77, 0.86] |

**Figure 3.**
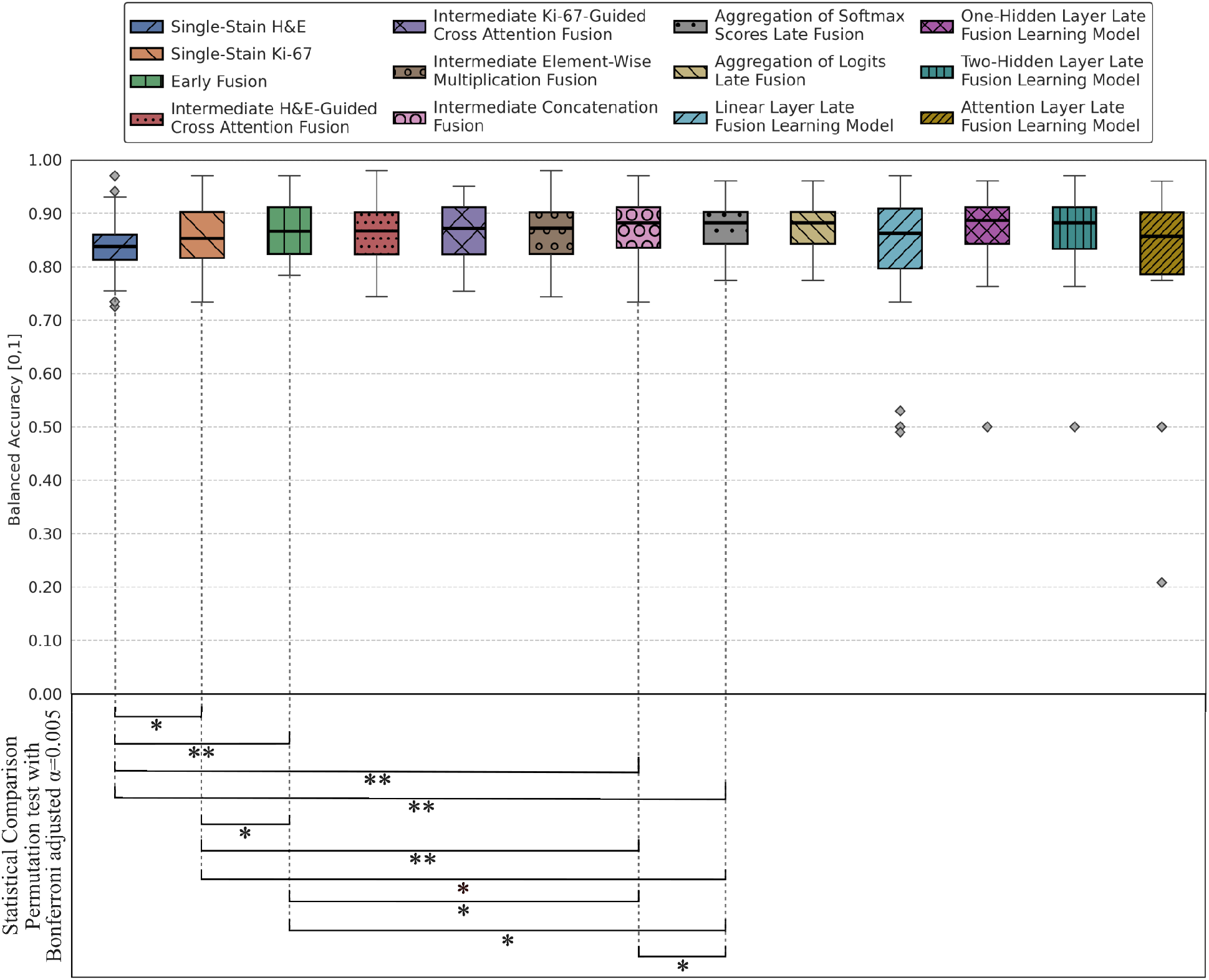
Boxplots summarizing balanced accuracy for LGG versus HGG classification on the test sets, computed across 50 non-parametric bootstrap replicates. Statistical comparisons are performed using a two-sided permutation test at a significance level of *α* = 0.05 between the single-stain models, early fusion, best-performing intermediate fusion (concatenation), and the best-performing late fusion (aggregation of softmax scores). Double asterisks (**) indicate statistically significant differences after Bonferroni correction, with an adjusted significance level of *α* = 0.05*/*10 = 0.005.

**Figure 4.**
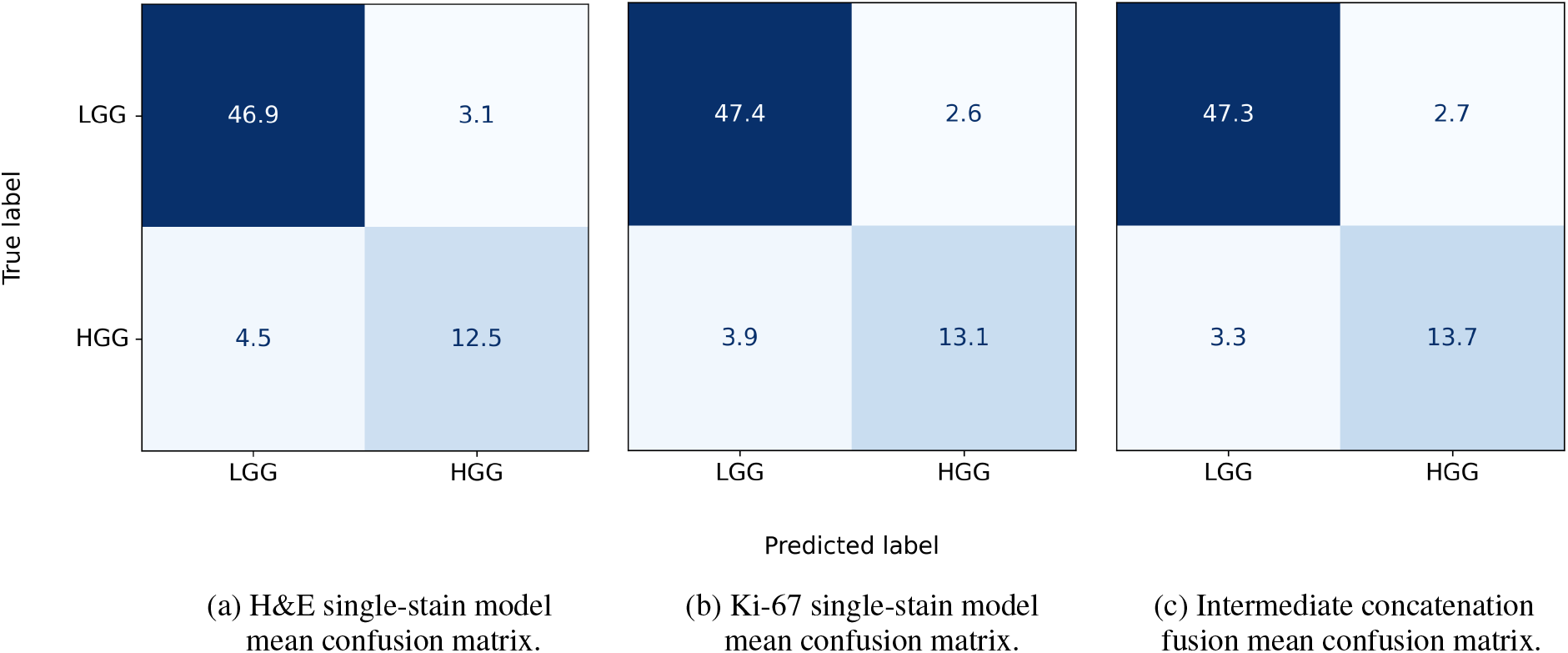
Mean confusion matrices summarizing the LGG versus HGG binary classification on the test sets, computed across 50 non-parametric bootstrap replicates, for (a) the H&E single-stain model, (b) the Ki-67 single-stain model, and (c) the intermediate concatenation fusion (best-performing fusion model).

**Table 3.**
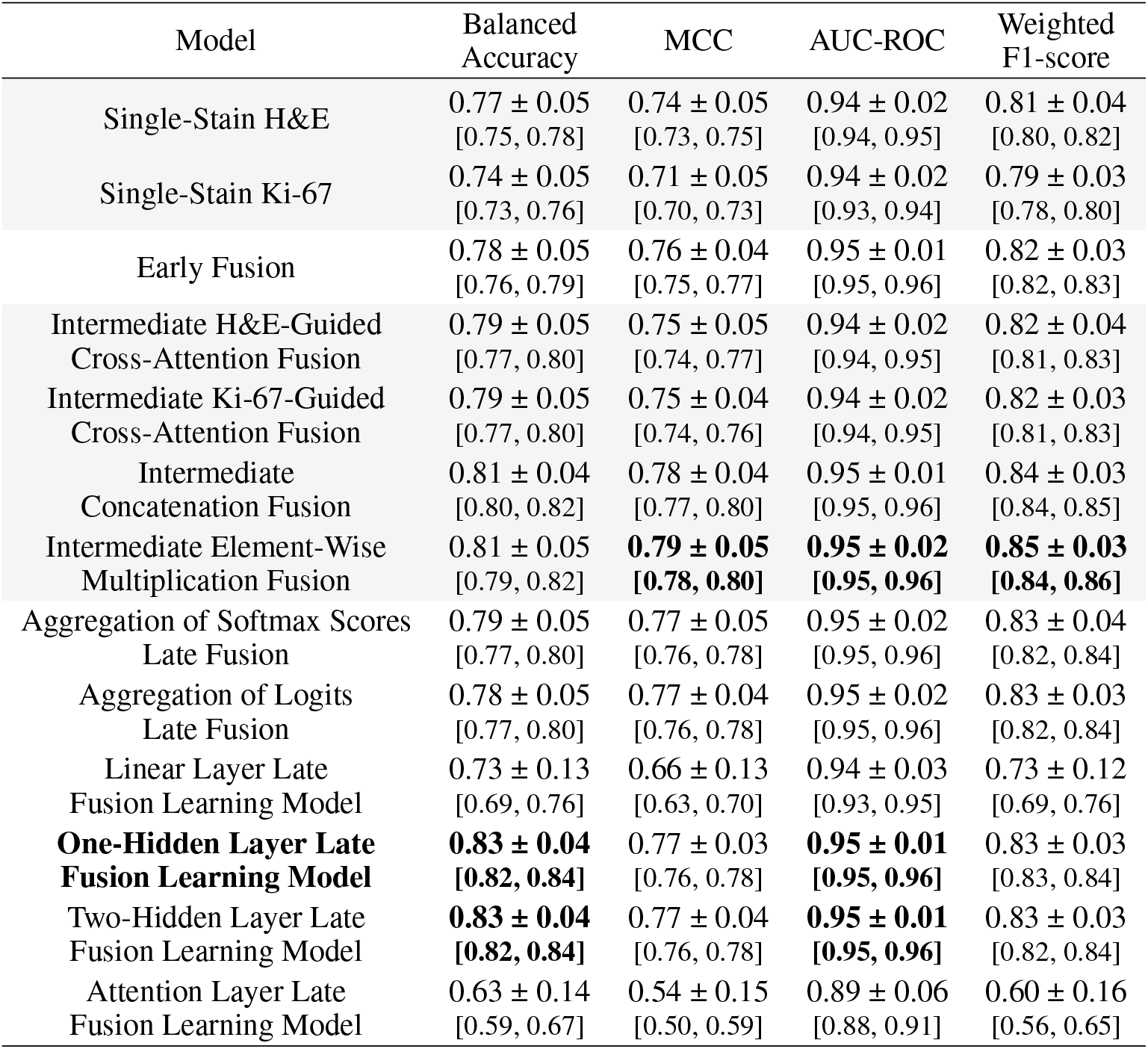
Classification performance between 5 tumor types on the test sets. Metrics are reported as a mean ± standard deviation with 95% confidence intervals (CI) shown in brackets, computed across 50 replicates of non-parametric bootstrapping. The best performing approach and metrics are highlighted in bold.

**Figure 5.**
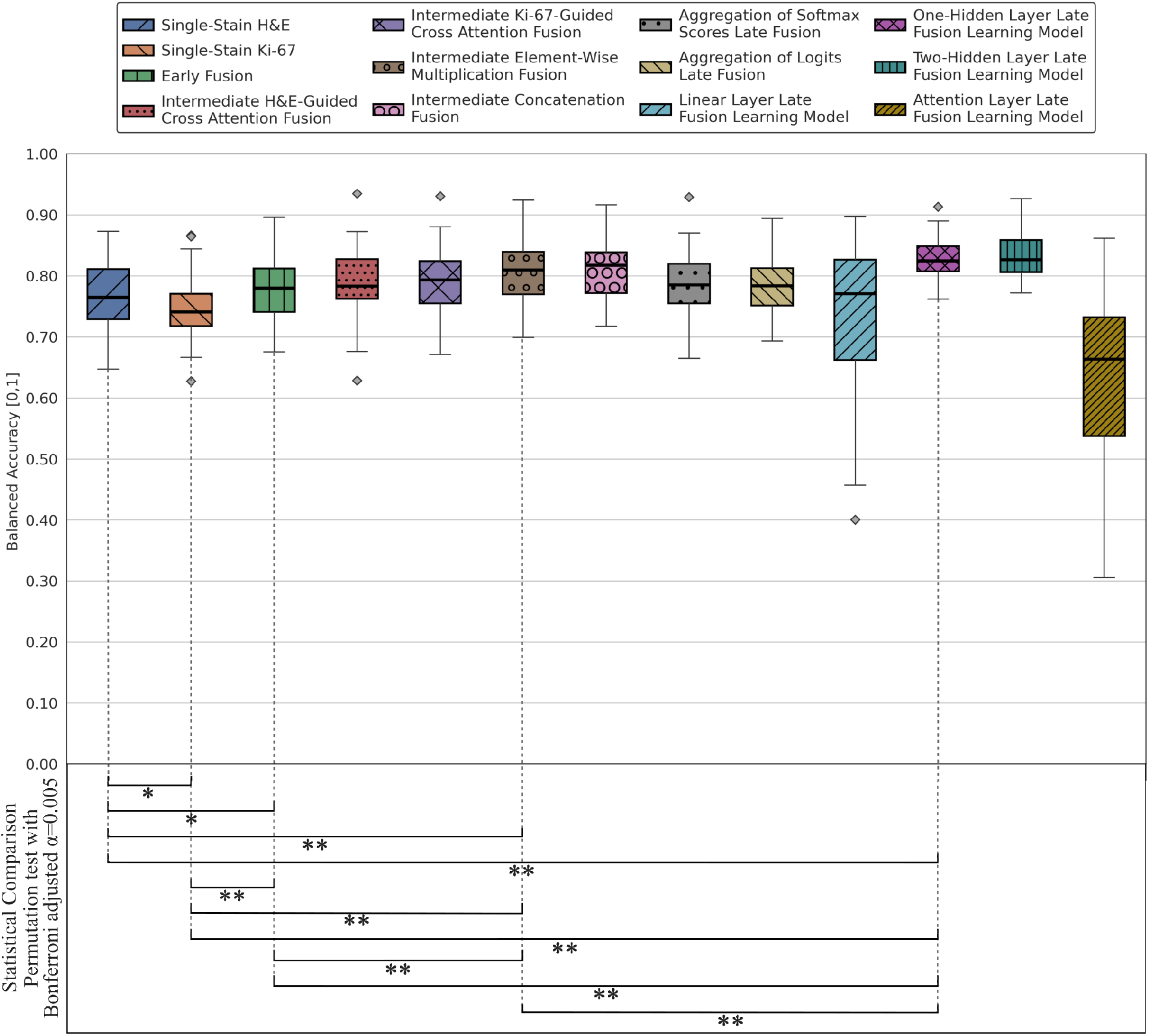
Boxplots summarizing balanced accuracy for the 5-class tumor type classification on the test sets, computed across 50 non-parametric bootstrap replicates. Statistical comparisons are performed using a two-sided permutation test at a significance level of *α* = 0.05 between the single-stain models, early fusion, best-performing intermediate fusion (element-wise multiplication), and the best-performing late fusion (one hidden layer learning model). Double asterisks (**) indicate statistically significant differences after Bonferroni correction, with an adjusted significance level of *α* = 0.05*/*10 = 0.005.

Most multi-stain experiments fusing H&E and Ki-67 demonstrated improved performance compared to the single-stain models. The early fusion, intermediate fusions, aggregation of softmax scores/logits late fusion, one-hidden layer and two-hidden layers late fusion learning models produced comparable results, with marginal differences in some metrics. The linear-layer and attention layer late fusion learning models performed worse than the single-stain models across all metrics. The intermediate concatenation fusion and aggregation of softmax scores were selected as the best-performing among the intermediate and late fusion approaches and included in the statistical comparisons.

The statistical test with Bonferroni-adjusted *α* = 0.005 showed that early fusion, intermediate concatenation fusion, and softmax score aggregation indicated significant differences over the H&E single-stain model across all metrics (*p <* 0.005). The comparison between the Ki 67 single-stain model and intermediate concatenation fusion demonstrated statistically significant differences in balanced accuracy, MCC, and weighted F1-score. The aggregation of softmax scores differed significantly from the Ki-67 model in MCC, AUC-ROC, and weighted F1- score. Moreover, the only significant difference between the early fusion and the Ki-67 single-stain model was in AUC-ROC. Between the fusion strategies, the only statistically significant difference was observed in AUC-ROC between early fusion and intermediate concatenation fusion. The intermediate concatenation fusion was selected as the best-performing approach because of its higher balanced accuracy.

In additional pairwise statistical comparisons conducted at a significance level of *α* = 0.05, aggregation of logits late fusion demonstrated statistically significant differences compared to the H&E and Ki-67 single-stain models in all metrics. In contrast, intermediate elementwise multiplication fusion, one-hidden layer and two-hidden layer late fusion learning models showed significant differences compared to the H&E single-stain model across all metrics, whereas no statistically significant differences were observed between these fusions and the Ki-67 single-stain model. The intermediate cross-attention fusions showed significant differences compared to the H&E single-stain model in all metrics except in the AUC-ROC, and only the intermediate H&E-guided cross-attention fusion showed significant differences compared to the Ki-67 single-stain model except in the AUC-ROC.

### 5.2 5-class Tumor Type Classification Performance

The results of the 5-class tumor type classification task are presented in Table 3 and Figures 5 and 6. The H&E single-stain model outperformed the Ki-67 single-stain model across most evaluation metrics, achieving higher balanced accuracy, MCC, and weighted F1-score, with both stains demonstrating the same AUC-ROC. The permutation test revealed significant differences in all metrics at a significance level of *α* = 0.05.

The fusion strategies generally outperformed the single-stain models. The one-hidden layer and two-hidden layers late fusion learning models achieved the highest balanced accuracy, with the one-hidden layer learning model selected as the best-performing late fusion approach because of its simplicity. The early fusion, intermediate crossattention fusions, the aggregation of softmax scores and logits late fusion provided comparable results to the H&E single-stain model and better performance compared to the Ki-67 single-stain model. The element-wise multiplication intermediate fusion, was selected as the best-performing intermediate fusion, with concatenation intermediate fusion showing similar performance. In contrast, the linear-layer and attention layer late fusion learning models showed worse results compared to the single-stain models.

The permutation test with Bonferroni-adjusted *α* = 0.005, showed statistically significant differences between the one-hidden layer late fusion learning model and the single-stain models in all metrics. Early fusion demonstrated significant differences compared to the H&E single-stain model in MCC, AUC-ROC, and weighted F1-score, and in all metrics compared to the Ki-67 single-stain model. Similarly, the element-wise multiplication intermediate fusion showed statistically significant differences compared to both single-stain models in all metrics. Statistical comparison among the fusion strategies revealed significant differences across most metrics, with AUC not differing significantly between the element-wise multiplication intermediate fusion and the one-hidden layer late fusion learning model, while early fusion and one-hidden layer late fusion learning model differed significantly only in balanced accuracy. The one-hidden layer late fusion learning model was selected as the best-performing approach because of its higher balanced accuracy.

In further pairwise comparisons conducted at a significance level of *α* = 0.05, the aggregation of softmax scores and logits outperformed the H&E and Ki-67 single-stain models, with significant differences in all metrics. The intermediate H&E-guided cross-attention fusion showed significant differences compared to the Ki-67 single-stain model across all metrics, and compared to the H&E single-stain model in all metrics except AUC-ROC. The intermediate Ki-67-guided cross-attention fusion differed significantly from the Ki-67 single-stain model in all metrics except AUC-ROC, while significant differences compared to the H&E single-stain model were observed in balanced accuracy and F1-score.

### 5.3 Interpretability and Explainability

The Figures 7, 8, 9 and 10 show examples of WSIs with their corresponding attention heatmaps and Ki-67 LI, negative and positive cell density maps. In the attention heatmaps, red and blue regions indicate high and low model attention, respectively. The HGG and LGG attention heatmaps were generated by the respective H&E and Ki-67 LGG vs HGG single-stain models, using the best-performing fold of the intermediate concatenation fusion model. Similarly, the ependymoma and medulloblastoma attention heatmaps were obtained from the respective H&E and Ki-67 5-class single-stain models, using the best-performing fold of the one-hidden-layer late fusion learning model. In the Ki-67 LI maps, the red and blue regions depict the proliferating and non-proliferating tumor cell ratios with the colorbar indicating the ratio of LI. In the negative and positive cell density maps, the color bar provides a scale of the density values corresponding to regions with high and low density of negative and positive cells.

**Figure 6.**
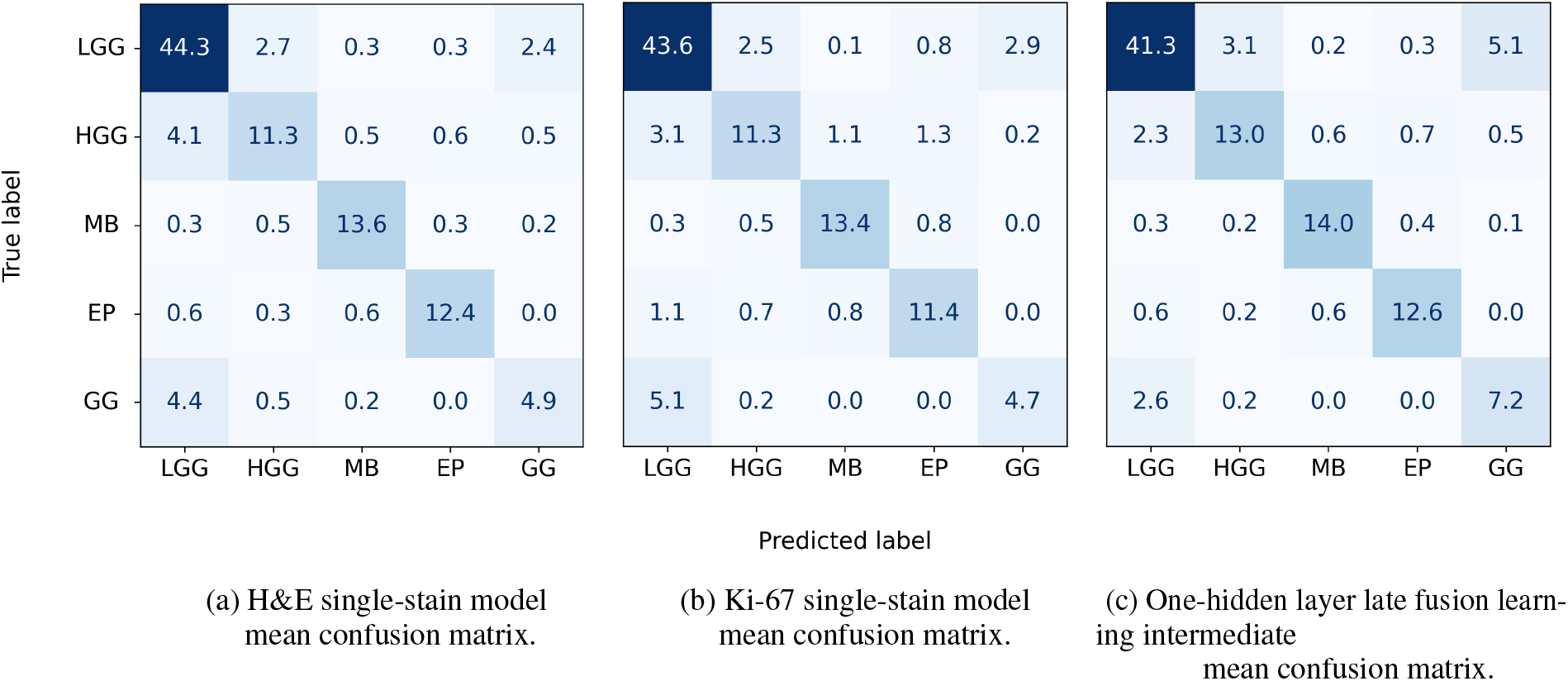
Mean confusion matrices summarizing the 5-class tumor type classification on the test sets, computed across 50 non-parametric bootstrap replicates, for (a) the H&E single-stain model, (b) the Ki-67 single-stain model, and (c) one-hidden layer late fusion learning model (best-performing fusion model).

**Figure 7.**
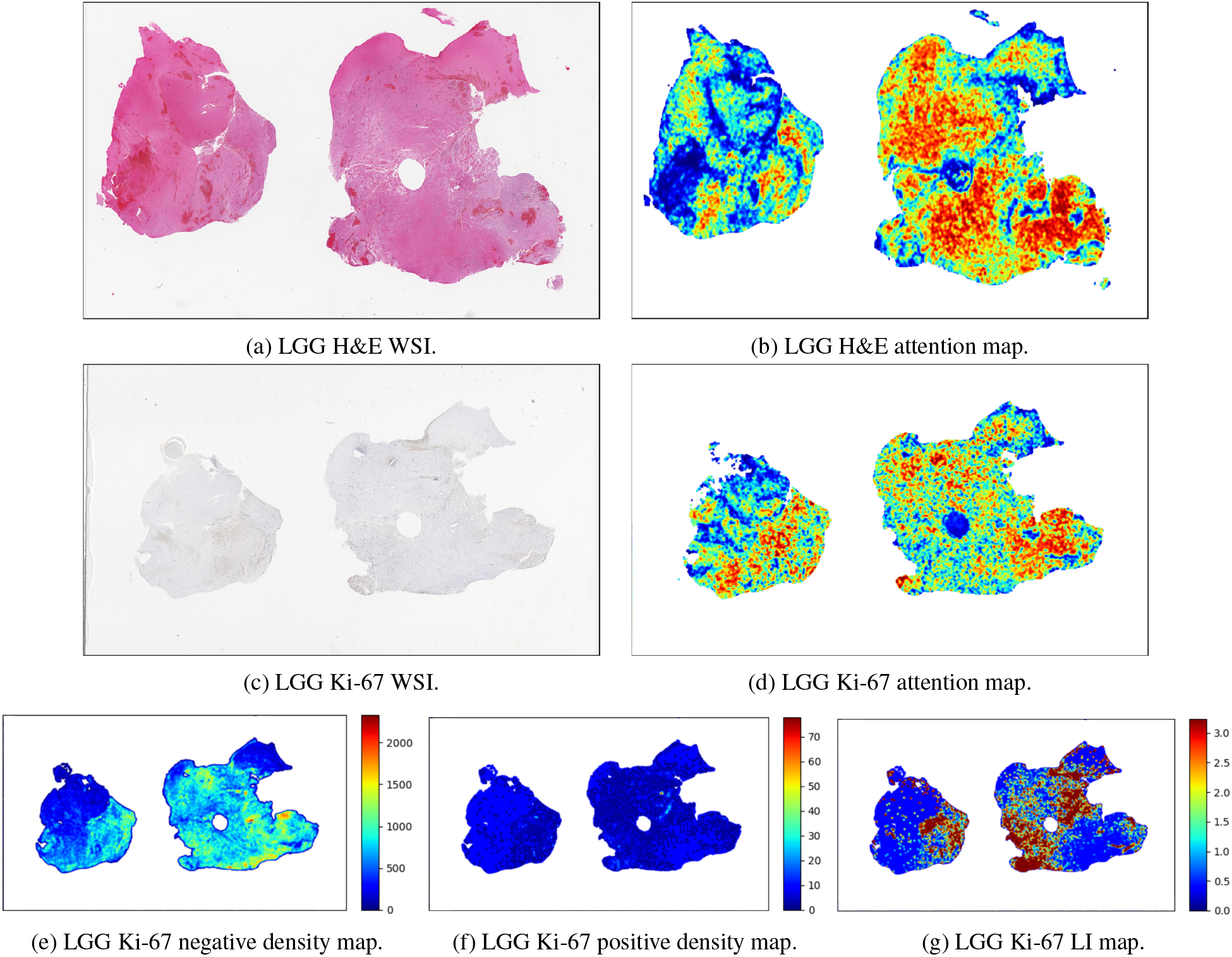
Representative LGG case showing (a) H&E WSI with its corresponding (b) attention map generated by the LGG vs HGG single-stain H&E model, (c) Ki-67 WSI with a LI of 3.24% with its corresponding (d) attention map generated by the LGG vs HGG single-stain Ki-67 model, (e) Ki-67 negative density map, (f) Ki-67 positive density map, and (g) Ki-67 LI map (*ρ* = 0.873 correlation with the heatmap).

**Figure 8.**
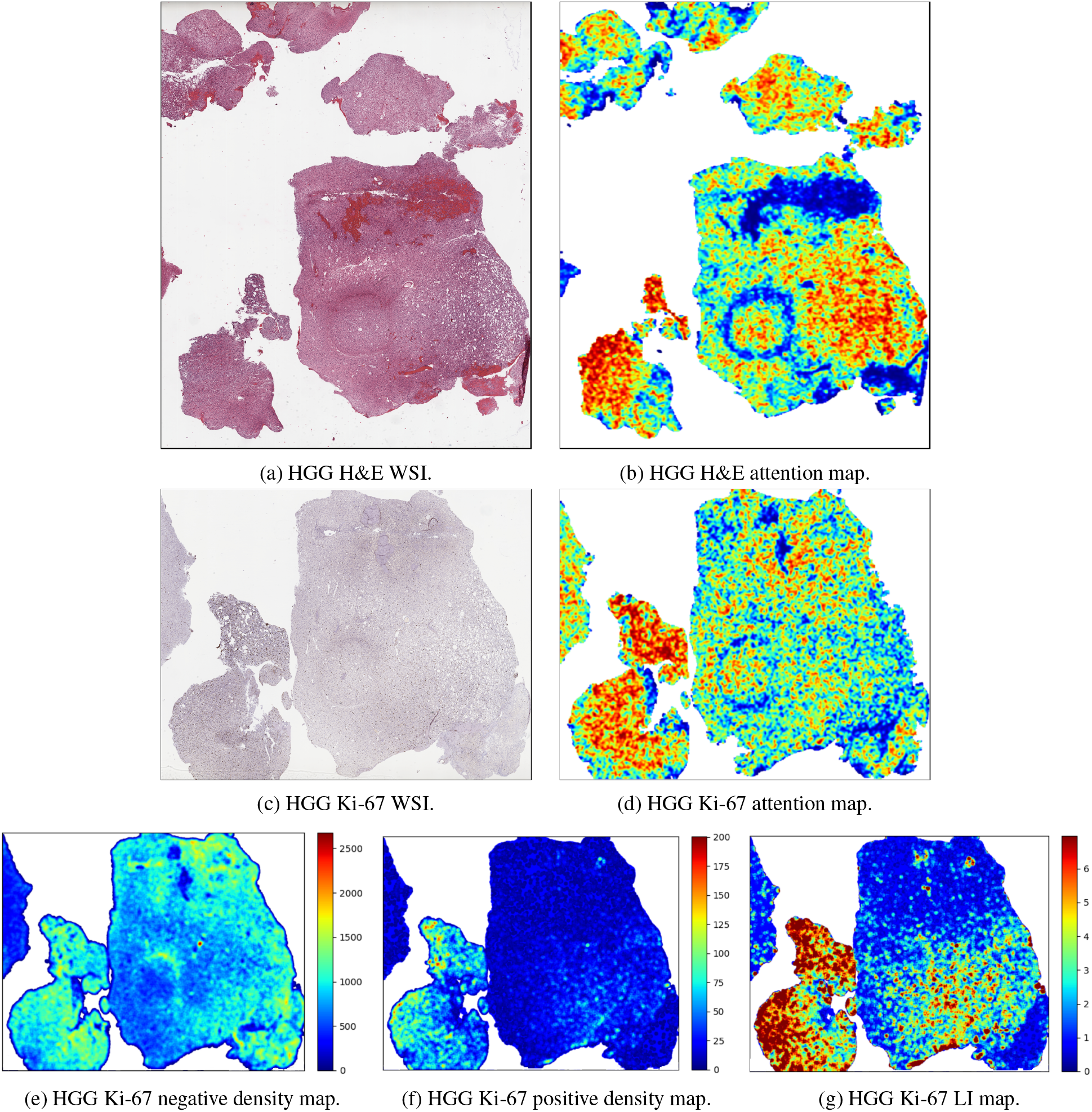
Representative HGG case showing (a) H&E WSI with its corresponding (b) attention map generated by the LGG vs HGG single-stain H&E model, and (c) Ki-67 WSI with a LI of 6.79% with its corresponding (d) attention map generated by the LGG vs HGG single-stain Ki-67 model and (e) Ki-67 negative density map, (f) Ki-67 positive density map, and (g) Ki-67 LI map (*ρ* = 0.515 correlation with the heatmap).

**Figure 9.**
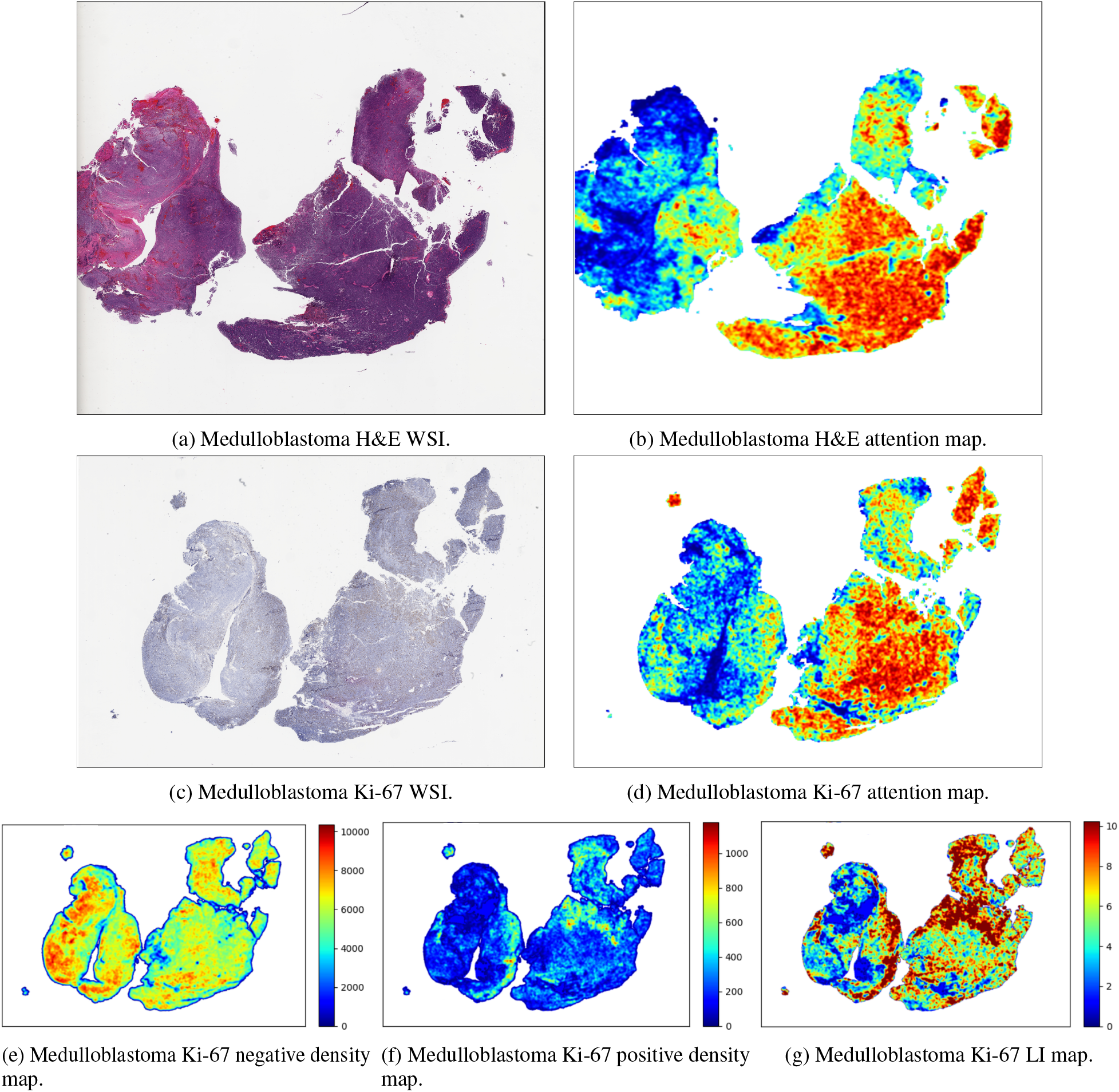
Representative medulloblastoma case showing (a) H&E WSI with its corresponding (b) attention map generated by the 5-class single-stain H&E model, (c) Ki-67 WSI with a LI of 10.23% with its corresponding (d) attention map generated by the 5-class single-stain Ki-67 model, (e) Ki-67 negative density map, (f) Ki-67 positive density map, and (g) Ki-67 LI map (*ρ* = 0.833 correlation with the heatmap).

**Figure 10.**
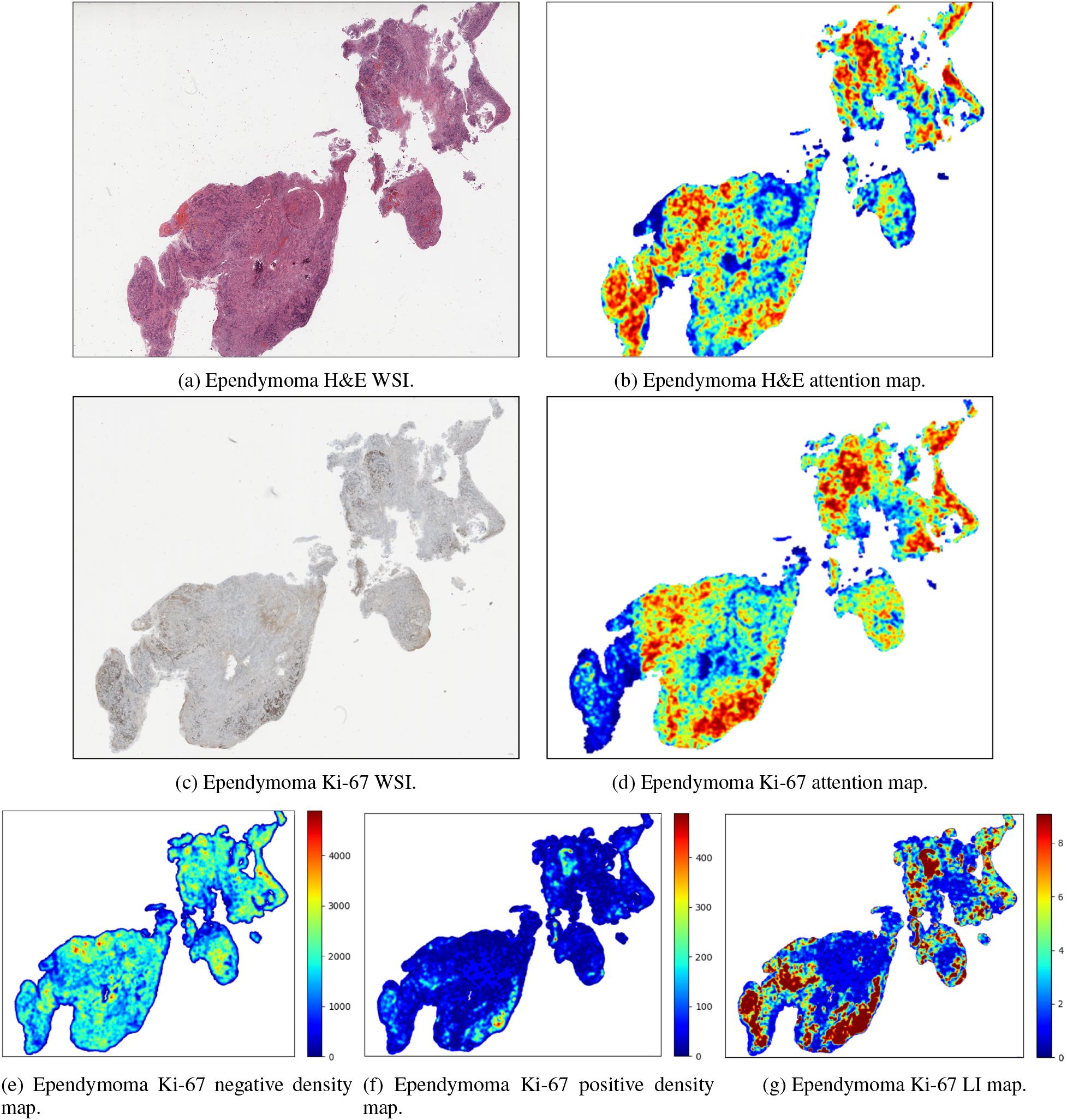
Representative ependymoma case showing (a) H&E WSI with its corresponding (b) attention map generated by the 5-class single-stain H&E model, (c) Ki-67 WSI with a LI of 9.09% with its corresponding (d) attention map generated by the 5-class single-stain Ki-67 model, (e) Ki-67 negative density map, (f) Ki-67 positive density map, and Ki-67 LI map (*ρ* = 0.844 correlation with the heatmap).

The Spearman’s rank correlation coefficients between the attention heatmaps and Ki-67 LI, negative and positive cell density maps, indicated a moderate to strong positive relationship across all tumor types as presented in Table 4. In the examples shown in the Figures 7, 8, 9 and 10, the visual inspection suggested that the Ki-67 attention heatmaps demonstrated overlaps with Ki-67 LI, negative and positive cell density maps.

## 6 Discussion

In this study, the classification of pediatric brain tumor families/types is investigated using deep learning on unregistered H&E and Ki-67 slides. Additionally, it was examined whether the fusion of the unregistered H&E and Ki-67 slides could improve the predictive performance compared to only using a single stain modality. The experiments indicated that the fusion approaches provided better and statistically significant different results compared to the individual stain models.

### 6.1 Classification Performance

In the binary classification task distinguishing LGG from HGG, single-stain models demonstrated strong performance, with the Ki-67 model outperforming the H&E model. These results are consistent with the clinical role of Ki-67 as a cellular proliferation marker, which is commonly used to distinguish glioma grades, suggesting that relevant features were captured by the Ki-67 single-stain model. Most fusion strategies outperformed the single-stain models, with intermediate concatenation fusion achieving the best predictive performance. The confusion matrices in Figure 4 showed that the intermediate concatenation fusion reduced the number of HGG misclassifications compared to both single-stain models, while achieving a similar number of correct LGG classifications to the Ki-67 single-stain model. Early fusion, intermediate element-wise multiplication fusion, intermediate crossattention fusion, aggregation of softmax scores and logits late fusion, and the one-hidden layer and two-hidden layers learning late fusion learning models provided results comparable to intermediate concatenation fusion.

In the five-class classification tumor type task, the H&E single-stain model outperformed the Ki-67 single-stain model. Most fusion strategies performed better and provided statistically significant different results compared to the single-stain models. The one-hidden layer late fusion learning model achieved the highest predictive performance and indicated statistically significant differences compared to the single-stain models and the fusion approaches. The confusion matrices depicted that the single-stain models and the one-hidden layer late fusion learning model accurately classified medulloblastoma and ependymoma, with the fusion approach achieving the best performance. Furthermore, the one-hidden layer late fusion learning model substantially improved the classification of ganglioglioma and HGG compared to the single-stain models, but performed worse in the LGG classification.

In both classification tasks, similar trends were observed in the performance of the fusion approaches. Early fusion outperformed the single-stain models, but did not provide the best results, since concatenating raw features at the patch-level prior to processing through CLAM might have limited the ability to capture stain-specific patterns. In contrast, intermediate and late fusions leveraged the most informative stain-specific features learned by the independently trained single-stain models, by utilizing patient-level attention vectors and CLAM outputs, respectively. The aggregation of softmax scores and logits late fusions provided comparable results, since the softmax scores are a normalized exponential transformation of the logits. Regarding the late fusion learning models, the performance of the one-hidden and two-hidden layer models suggested that moderate model complexity is sufficient to capture relationships between the two stain modalities. The lower performance of the linear and attention layer late fusion learning models may reflect the inability to capture nonlinear relationships, and a possible requirement for more data to effectively learn meaningful attention weights, respectively.

Overall, the experiments demonstrated that fusing H&E and Ki-67 images improved the predictive performance compared to single-stain models, supporting the hypothesis that the two stains provide complementary information. No specific fusion approach performed the best across both classification tasks, the optimal fusion approach might be related to task complexity, since the statistical comparisons indicated that the significant differences were more evident in the 5-class tumor type classification task compared to the binary grade classification task.

### 6.2 Interpretability and Explainability

The Spearman’s rank correlation coefficients *ρ* between the Ki-67 attention maps and the corresponding Ki-67 LI, negative and positive cell density maps demonstrated a moderate to strong positive correlation. Quantitative and qualitative analyses were limited to the Ki-67 attention heatmaps, since the LI, positive and negative cell density maps were the only reference for evaluation in this study, although additional features may also be captured by the Ki-67 single-stain model. The explainability of the H&E attention heatmaps was not investigated, as establishing ground truth would require manual annotation by a pathologist to label a broader spectrum of morphological features, such as cellularity, necrosis, and vascular patterns, to assess whether the H&E single-stain model captures these relevant features [26].

### 6.3 Challenges and Limitations

The primary limitation in this study is that conventional intermediate fusion approaches could not be implemented since the features were extracted using the pre-trained CONCHv1_5 histology foundation model, limiting the potential to fully share complementary information. Moreover, pre-training or fine-tuning the models specifically for pediatric tumors would be preferable, but the lack of extensive pediatric datasets is a challenge and represents a direction for future work. Moreover, datasets should include not only H&E but additional IHC and other stained WSIs, thereby providing the opportunity to extract further clinical value using deep learning. Regarding the dataset, one limitation is that tumor entities were excluded from the analysis given the incompatibility with the 2021 WHO classification guidelines for CNS tumors. More detailed tumor grading information, such as ependymoma grades, molecular subgroups of medulloblastoma, and slide-level diagnoses, would have enabled to conduct a more granular analysis and the investigation of potential performance improvements in clinical classification. Furthermore, the limited availability of additional stain modalities in sufficient numbers constrained the potential for exploring how information across multiple stains could be leveraged to improve predictive performance.

**Table 4.** Spearman’s rank correlation coefficients between Ki-67 attention heatmaps and LI maps, negative cell density maps, and positive cell density maps for LGG vs HGG and 5-class classification tasks.

| Classification Task | Tumor | Ki-67 LI map<br>Spearman’s $\rho$ | Negative Density Map<br>Spearman’s $\rho$ | Positive Density Map<br>Spearman’s $\rho$ |
| --- | --- | --- | --- | --- |
| LGG vs HGG | LGG | 0.797 | 0.742 | 0.814 |
|  | HGG | 0.651 | 0.610 | 0.641 |
|  | Overall | 0.761 | 0.710 | 0.767 |
| 5-class | LGG | 0.788 | 0.723 | 0.803 |
|  | HGG | 0.615 | 0.581 | 0.576 |
|  | Medulloblastoma | 0.716 | 0.646 | 0.683 |
|  | Ependymoma | 0.714 | 0.645 | 0.701 |
|  | Ganglioglioma | 0.818 | 0.729 | 0.823 |
|  | Overall | 0.748 | 0.685 | 0.739 |

## 7 Conclusion

In this study, attention-based MIL and fusion approaches were implemented on features extracted from unregistered H&E and Ki-67 WSIs using a histology foundation model to classify pediatric brain tumors. Most of the fusion approaches of the H&E and Ki-67 WSIs provided improved predictive performance compared to the single-stain models, indicating that the two stain modalities provide complementary information. The correlation analysis and visual comparison between the attention heatmaps and the Ki-67 LI, negative and positive cell density maps suggest that these features contribute to the model’s predictions, in addition to other unassessed features. Overall, these findings support multi-stain fusion as a promising approach for improving pediatric brain tumor diagnosis, with potential for further improvement through the fusion of additional modalities, such as molecular data, and application in other cancer types.

## Acknowledgements and Funding

The research was made possible in part due to The Children’s Brain Tumor Tissue Consortium (CBTTC)/ The Children’s Brain Tumor Network (CBTN). The study was financed by Swedish Childhood Cancer Foundation (MT2021-0011, MT2022-0013), Joanna Cocozza’s Foundation (2025-2026), Linköping University’s Cancer Strength Area (2024), Medical Research Council of Southeast Sweden (FORSS-1011571). Lindblad was supported by the Swedish Cancer Society (25 4859 Pj).

## Data and Code Availability

The open-access dataset used in this study was obtained from the CBTN (https://cbtn.org). The detailed instructions, python environment and scripts for reproducing the experiments are available in the associated GitHub repository https://github.com/Christoforos-Spyretos/CBTN_Histology_Multi_Stain. The weights for CONCHv1_5 pre-trained model are available at https://github.com/mahmoodlab/CONCH?tab=readme-ov-file and the original code for CLAM is available at https://github.com/mahmoodlab/CLAM.

## Ethical Approval Statement

Not applicable. The dataset is anonymous and openaccess provided by CBTN.

## References

[1] J. Ferlay et al. Global Cancer Observatory: Cancer Today. Lyon, France: International Agency for Research on Cancer. https://gco.iarc.who.int/today. xAccessed: 2025. 2025.

[2] David N Louis et al. “The 2021 WHO classification of tumors of the central nervous system: a summary”. In: Neuro-oncology 23.8 (2021), pp. 1231–1251.

[3] Andrey Bychkov and Michael Schubert. “Constant Demand, Patchy Supply”. In: 88 (Feb. 2023), pp. 18–27.

[4] Massimo Salvi et al. “The impact of pre-and post-image processing techniques on deep learning frameworks: A comprehensive review for digital pathology image analysis”. In: Computers in Biology and Medicine 128 (2021), p. 104129.

[5] Andrew H Song et al. “Artificial intelligence for digital and computational pathology”. In: Nature Reviews Bioengineering 1.12 (2023), pp. 930–949.

[6] Jan-Philipp Redlich et al. “Applications of artificial intelligence in the analysis of histopathology images of gliomas: a review”. In: npj Imaging 2.1 (2024), p. 16.

[7] Marc-André Carbonneau et al. “Multiple instance learning: A survey of problem characteristics and applications”. In: Pattern recognition 77 (2018), pp. 329–353.

[8] Michael Gadermayr and Maximilian Tschuchnig. “Multiple instance learning for digital pathology: A review of the state-of-the-art, limitations & future potential”. In: Computerized Medical Imaging and Graphics 112 (2024), p. 102337.

[9] Maximilian Ilse, Jakub M Tomczak, and Max Welling. “Deep multiple instance learning for digital histopathology”. In: Handbook of Medical Image Computing and Computer Assisted Intervention. Elsevier, 2020, pp. 521–546.

[10] Shengjia Chen et al. “Benchmarking embedding aggregation methods in computational pathology: A clinical data perspective”. In: arXiv preprint arXiv:2407.07841 (2024).

[11] Daniel Shao et al. “Do Multiple Instance Learning Models Transfer?” In: Forty-second International Conference on Machine Learning.

[12] Jonas Dippel et al. “RudolfV: a foundation model by pathologists for pathologists”. In: arXiv preprint arXiv:2401.04079 (2024).

[13] Julián N Acosta et al. “Multimodal biomedical AI”. In: Nature medicine 28.9 (2022), pp. 1773–1784.

[14] Adrienne Kline et al. “Multimodal machine learning in precision health: A scoping review”. In: NPJ digital medicine 5.1 (2022), p. 171.

[15] Angela N Viaene. “Pediatric brain tumors: A neuropathologist’s approach to the integrated diagnosis”. In: Frontiers in Pediatrics 11 (2023), p. 1143363.

[16] Antonio d’Amati et al. “Pediatric CNS tumors and 2021 WHO classification: what do oncologists need from pathologists?” In: Frontiers in molecular neuroscience 17 (2024), p. 1268038.

[17] Lian Tao Li et al. “Ki67 is a promising molecular target in the diagnosis of cancer”. In: Molecular medicine reports 11.3 (2015), pp. 1566–1572.

[18] Sigrid Uxa et al. “Ki-67 gene expression”. In: Cell Death & Differentiation 28.12 (2021), pp. 3357–3370.

[19] Rikke H Dahlrot et al. “Prognostic role of Ki-67 in glioblastomas excluding contribution from non-neoplastic cells”. In: Scientific reports 11.1 (2021), p. 17918.

[20] Daniele Armocida et al. “Is Ki-67 index overexpression in IDH wild type glioblastoma a predictor of shorter Progression Free survival? A clinical and Molecular analytic investigation”. In: Clinical neurology and neurosurgery 198 (2020), p. 106126.

[21] Fu Zhao et al. “Prognostic value of Ki-67 index in adult medulloblastoma after accounting for molecular subgroup: a retrospective clinical and molecular analysis”. In: Journal of Neuro-Oncology 139.2 (2018), pp. 333–340.

[22] Vikas Sharma et al. “P53 and Ki-67 expression in primary pediatric brain tumors: Does it correlate with presentation, histological grade, and outcome?” In: Asian journal of neurosurgery 13.04 (2018), pp. 1026–1032.

[23] Subhalakshmi Sengupta et al. “A study of histopathological spectrum and expression of Ki-67, TP53 in primary brain tumors of pediatric age group”. In: Indian Journal of Medical and Paediatric Oncology 33.01 (2012), pp. 25–31.

[24] Eun-Ik Son et al. “Immunohistochemical analysis for histopathological subtypes in pediatric medulloblastomas”. In: Pathology international 53.2 (2003), pp. 67–73.

[25] Sandra Steyaert et al. “Multimodal deep learning to predict prognosis in adult and pediatric brain tumors”. In: Communications Medicine 3.1 (2023), p. 44.

[26] Iulian Emil Tampu et al. “Pediatric brain tumor classification using digital pathology and deep learning: Evaluation of SOTA methods on a multi-center Swedish cohort”. In: Brain Pathology (2025), e70029.

[27] Omneya Attallah and Shaza Zaghlool. “AI-based pipeline for classifying pediatric medulloblastoma using histopathological and textural images”. In: Life 12.2 (2022), p. 232.

[28] Marcel Bengs et al. “Medulloblastoma tumor classification using deep transfer learning with multi-scale EfficientNets”. In: Medical Imaging 2021: Digital Pathology. Vol. 11603. SPIE. 2021, pp. 70–75.

[29] Jon Whitney et al. “Quantitative nuclear histomorphometry predicts molecular subtype and clinical outcome in medulloblastomas: Preliminary findings”. In: Journal of Pathology Informatics 13 (2022), p. 100090.

[30] Guillaume Jaume et al. “Multistain pretraining for slide representation learning in pathology”. In: European Conference on Computer Vision. Springer. 2024, pp. 19–37.

[31] Shengyi Hua et al. “PathoDuet: foundation models for pathological slide analysis of H&E and IHC stains”. In: Medical Image Analysis 97 (2024), p. 103289.

[32] Joshua A Shapiro et al. “OpenPBTA: The Open Pediatric Brain Tumor Atlas”. In: Cell genomics 3.7 (2023).

[33] Jena V Lilly et al. “The children’s brain tumor network (CBTN)-Accelerating research in pediatric central nervous system tumors through collaboration and open science”. In: Neoplasia 35 (2023), p. 100846.

[34] Christoforos Spyretos et al. “Early fusion of H&E and IHC histology images for pediatric brain tumor classification”. In: MICCAI Workshop on Computational Pathology with Multimodal Data (COMPAYL).

[35] Christoforos Spyretos et al. “Quantification of Ki-67 labeling index in pediatric brain tumor immunohistochemistry images”. In: Journal of Neuropathology & Experimental Neurology (2026), laf163.

[36] Ming Y Lu et al. “Data-efficient and weakly supervised computational pathology on whole-slide images”. In: Nature biomedical engineering 5.6 (2021), pp. 555–570.

[37] Tong Ding et al. “A multimodal whole-slide foundation model for pathology”. In: Nature Medicine (2025), pp. 1–13.

[38] Richard J Chen et al. “Towards a general-purpose foundation model for computational pathology”. In: Nature medicine 30.3 (2024), pp. 850–862.

[39] Ming Y Lu et al. “A visual-language foundation model for computational pathology”. In: Nature medicine 30.3 (2024), pp. 863–874.

[40] Jana Lipkova et al. “Artificial intelligence for multimodal data integration in oncology”. In: Cancer cell 40.10 (2022), pp. 1095–1110.

[41] Ashish Vaswani et al. “Attention is all you need”. In: Advances in neural information processing systems 30 (2017).

[42] Ngoc-Quang Nguyen et al. “MulinforCPI: enhancing precision of compound–protein interaction prediction through novel perspectives on multi-level information integration”. In: Briefings in Bioinformatics 25.1 (2024), bbad484.

[43] Nicole Bussola et al. “AI slipping on tiles: data leakage in digital pathology”. In: International Conference on Pattern Recognition. Springer. 2021, pp. 167–182.

